# A specific Ret receptor isoform is required for pioneer axon outgrowth and growth cone dynamics

**DOI:** 10.1101/424432

**Authors:** Adam M. Tuttle, Catherine M. Drerup, Molly H. Marra, Alex V. Nechiporuk

## Abstract

In many cases, axon growth and guidance are driven by pioneer axons, the first axons to grow in a particular region. Despite their dynamic pathfinding capabilities and developmental importance, there are very few pioneer neuron specific markers and thus their *in vivo* identification and functional interrogation have been difficult. We found that a Ret receptor isoform, Ret51, is highly enriched in peripheral sensory pioneer neurons and is required for pioneer axon outgrowth. *Ret* null mutant pioneer neurons differentiate normally; however, they displayed defects in growth cone morphology and formation of filopodia before pioneer axon extension prematurely halts. We also demonstrate loss-of-function of a retrograde cargo adaptor, JNK-interacting protein 3 (Jip3), phenocopied many of these axonal defects. We further found that loss of Jip3 led to accumulation of activated Ret receptor in pioneer growth cones, indicating a failure in the clearance of activated Ret from growth cones. Using an axon sever approach as well as *in vivo* analysis of axonal transport, we showed Jip3 specifically mediates retrograde, but not anterograde, transport of activated Ret51. Finally, live imaging revealed that Jip3 and Ret51 were retrogradely co-transported in pioneer axons, suggesting Jip3 functions as an adapter for retrograde transport of Ret51. Taken together, these results identify Ret51 as a molecular marker of pioneer neurons and elucidate an important isoform-specific role for Ret51 in axon growth and growth cone dynamics during development.

## Introduction

Pioneer neurons are the first to extend axons to a particular region or target, acting as a guide and scaffold for “follower” axons. Pioneer neurons are important in the developing CNS and PNS for the initial navigation to appropriate targets, proper follower axon pathfinding, and promoting follower axon outgrowth. Despite their unique role in neurodevelopment, the molecular program that specifies pioneer neuron identity and regulates their behavior is not well known. Recent studies describe multiple factors, including TAG-1^1^, FMI-1^2^, and AEX-3^3^, that promote pioneer axon growth and pioneer/follower interactions in mouse and *C. elegans* models. However, these are not exclusive pioneer markers because they are also expressed in follower neurons, too transient to use as a reliable marker for isolating cells or have additional roles in other neuronal subtypes during development. While some progress continues to be made in terms of what factors contribute to pioneer neuron identity and promote some unique axonal properties, specific molecular markers are sorely lacking and act as a barrier to advancing our fundamental understanding of pioneer neuron biology.

In the process of pathfinding, pioneer neuron growth cones must migrate forward, turn, and interact with the local ECM and signaling environment. Dynamic changes in actin cytoskeleton structure in the growth cone are critical for all of these processes. In particular, formation and motility of filopodial protrusions are critical for sensing extrinsic cues for growth cone guidance and are required for proper axon pathfinding and growth cone advancement^4,5^. One major regulator of axon outgrowth and pathfinding, both during development and following axonal injury, is signaling through neurotrophic factors, including NGF, BDNF, and GDNF. Of these, the best studied neurotrophins are Nerve Growth Factor (NGF) and Brain-derived Growth Factor (BDNF) that signal through the neurotrophin receptors TrkA and TrkB, respectively. NGF and BDNF can have effects locally on axon terminals or signal retrogradely from the axon terminal to the cell body and induce changes in gene expression. For example, NGF and BDNF can induce localized actin polymerization within growth cones, promoting growth cone protrusions through formation of finger-like F-actin bundles, filopodia, or promoting local growth cone “spreading” through formation of a sheet-like actin mesh network, lamellipodia^6-9^, which is required for growth cone steering and advancement. In contrast, NGF/BDNF long-range retrograde signaling is required for survival of sympathetic neurons, where retrograde transport of active endosomal signaling complexes promotes transcription of pro-survival/anti-apoptotic factors^10-12^. In addition to survival, retrograde transport of NGF/TrkA promotes sensory axon outgrowth by inducing transcriptional changes in Serum Response Factor^13^, whereas retrograde axonal transport of BDNF/TrkB is required for BDNF-mediated dendritic growth in cortical neurons^14^. In general, however, the transcriptional targets downstream of retrograde neurotrophin signaling that promote axon growth are still largely unknown.

The neurotrophin receptor Ret and its major ligand, Glial cell line-Derived Neurotrophic Factor (GDNF), have established roles in axon growth, enteric nervous system and kidney development, and cancer progression. Ret receptor has two primary expressed isoforms, Ret9 and Ret51, each with a unique amino acid sequence of the intracellular C-terminus. The two isoforms also have different signaling properties and roles during development *in vivo*. Ret9 is required for proper enteric neuron survival, migration, and differentiation as well as kidney development, while Ret51 is largely dispensable for these processes. Furthermore, monoisoformic RET9 mice (lacking RET51) are viable, while monoisoformic RET51 mice (lacking RET9) die as neonates^15^. These isoforms are also differentially trafficked and have differential susceptibility to degradation in both neurons and non-neuronal cells^16,17^. In sympathetic neuron culture, RET9 degrades more slowly than RET51, which is thought to allow retrograde trafficking of RET9 and ultimately promote neuronal survival^16^. In human epithelial cell lines, RET9 is transcribed at much higher levels than RET51, but RET51 is comparatively more frequently localized at the plasma membrane, recycled to the plasma membrane after endocytosis, and more rapidly internalized into endosomes after GDNF treatment^17,18^. Thus, Ret9 and Ret51 have notably different properties and responses to ligand binding even within the same cell.

Ret/GDNF signaling promotes sympathetic^19^, sensory^20^, and motor^21^ axon growth and pathfinding. Long-range retrograde RET/GDNF trafficking from distal axons is required for survival of sensory (DRG) neurons and older (15 days *in vitro*), but not younger (10 days *in vitro*), sympathetic neurons in rat primary cell cultures^16,22^. In sympathetic neurons, long-range retrograde transport of GDNF ligand from axon terminals to cell bodies has been observed^22,23^ but, thus far, retrograde transport of GDNF/Ret in sensory neurons has not. Ret/GDNF signaling can also promote sympathetic axon growth *in vitro*, however, retrograde signaling/transport is not required for this role^16^. Long-range signaling through GDNF/Ret is required for motor neuron extension in developing mouse embryos where loss of GDNF ligand expression in target muscles leads to reduced neuronal expression of *Pea3*, a transcription factor required for proper cell body positioning and dendrite patterning in the neurons whose axons receive GDNF^24,25^. However, the mechanisms mediating Ret/GDNF retrograde transport and the full extent of the Ret/GDNF downstream transcriptional response mediating axon growth are unknown and *in vivo* studies of Ret retrograde transport and signaling are sorely lacking^26^.

*In vivo* study of long range neurotrophin signaling, receptor transport, and pioneer axon growth is technically challenging. Zebrafish embryos offer a number of advantages for these studies, including optical accessibility, rapid embryonic development, and amenability to transgenesis and genetic manipulation. In this study, we use the long sensory axons of the zebrafish posterior lateral line ganglion (pLLg) neurons because their planar character, superficial localization, and relatively rapid pioneer axon outgrowth make them uniquely suited for live imaging. The pLL senses water movement through mechanical stimulation of sensory organs (neuromasts) that transduce this input through the pLL axons innervating them. During extension, 3 to 6 pLL pioneer axons are guided by the pLL primordium (pLLp), a group of cells that migrate along the trunk from 22 to 48 hours post-fertilization (hpf). Pioneer growth cones typically have longer and more elaborate filopodial protrusions compared to followers and this is the case in pLLg pioneer growth cones, which associate extensively with the migrating pLLp^27^. The pLL primordium expresses GDNF and this ligand production is required for proper pLLg axon outgrowth^28^, but not pLLg neuron specification or survival^28^. This allowed us to explore the specific mechanisms of long-range Ret-mediated axon outgrowth in sensory pioneer neurons.

We find that Ret is specifically expressed in pioneer axons and *ret51*, but not *ret9*, isoform is required for pioneer axon outgrowth. In the absence of Ret, pioneer axons display reduced growth cone volume and fewer filopodia, which may underlie the observed pioneer axon outgrowth defects. We also show that JNK-interacting protein 3 (Jip3) is required for retrograde transport of activated Ret51 from pioneer growth cones. In *jip3* mutants, retrograde transport of Ret51 is reduced, leading to accumulation of activated Ret in pioneer growth cones. Finally, we observe live retrograde, but not anterograde, co-transport of tagged Ret51 and Jip3 in pioneer sensory axons. This work describes a novel requirement for Ret51 in axon growth and pioneer neuron development and identifies a novel regulator of phosphorylated Ret51 retrograde transport.

## Results

### Ret expression is elevated in pioneer neurons during axon extension

Previous studies showed that *ret* is expressed in the pLLg neurons and the Ret ligand GDNF, which is expressed by the pLLp, is required pLLg axon extension^28^. Because Ret receptor isoforms Ret9 and Ret51 have distinct properties, we asked which isoform is required for sensory axon extension. We detected expression of both *ret9* and *ret51* in the pLLg during pioneer axon outgrowth (30 hours post-fertilization – hpf) by *in situ* hybridization. *ret9* is expressed broadly throughout the pLLg (Fig. 1A), while *ret51* is highly expressed in a subset of the pLLg in the dorsal/anterior region (Fig.1B).

**Figure 1:**
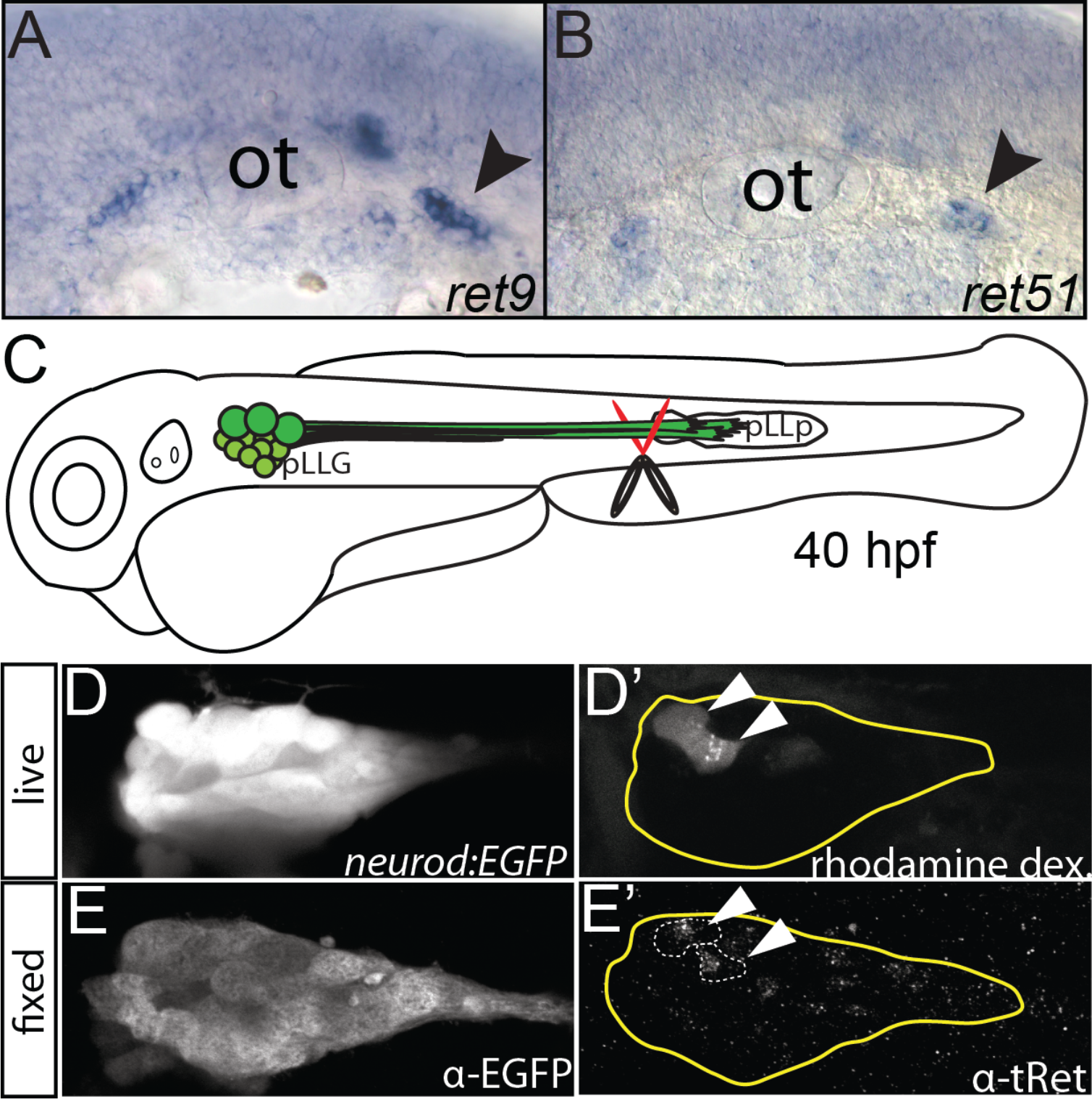
*ret51* is highly expressed in pLL pioneer neurons. (A,B) Lateral view, *in situ* hybridization reveals
*ret9* and *ret51* are both expresed during axon extension (30 hpf) in the pLLg. Note that, *ret51* is expressed in a small subset of pLLg cells in the more dorsal region (arrowhead =pLLg; ot=otic vesicle). (C) Diagram of pLL nerve sever experiment. Distal pioneer axons of TgBAC(*neurod:EGFP)* transgenic embryos were severed during extension using rhodamine dextran soaked scissors. pLLp = posterior lateral line primordium. Live image of EGFP (D) and rhodamine labeled pLLg pioneer neuron cell bodies (D’) 3 hours post-sever. Immunostaining for EGFP (E) and α-total Ret (E’, tRet) shows enriched Ret protein present in the rhodamine positive cells (arrowheads) (8 of 10 cells analyzed from n=5 embryos).

Based on elevated levels of *ret51* mRNA expression in the dorsal region of the ganglion, which was previously attributed to the location of pioneer neurons^29^, we tested whether Ret protein was enriched in extending pioneer neurons. To label cell bodies of the extending pioneer pLLg axons, tails of 40 hpf *TgBAC(neurod:EGFP)^nl1^* transgenic zebrafish embryos (hereafter referred to as *neurod:EGFP*)^30^ were clipped just proximal to the pioneer axon terminals with rhodamine dextran-soaked scissors (Fig. 1C). After 3 hours, embryos were imaged live to note the location of rhodamine-labeled cell bodies in pLLg, then fixed and immunostained with α-Ret primary antibody (rabbit, for α-Ret antibody validation, see Suppl. Fig. 2) and α-rabbit AlexaFluor 647 secondary (Thermo Fisher) to distinguish α-Ret antibody signal from rhodamine. We found that the majority of rhodamine dextran labeled pioneer axons also displayed high levels of Ret protein (Fig. 1D’,E’). The dorsal localization of the cell bodies positive for Ret and enrichment of *ret51* transcript in the same location of the ganglion strongly suggest that Ret51 receptor isoform is specifically expressed in pioneer neurons.

**Figure 2:**
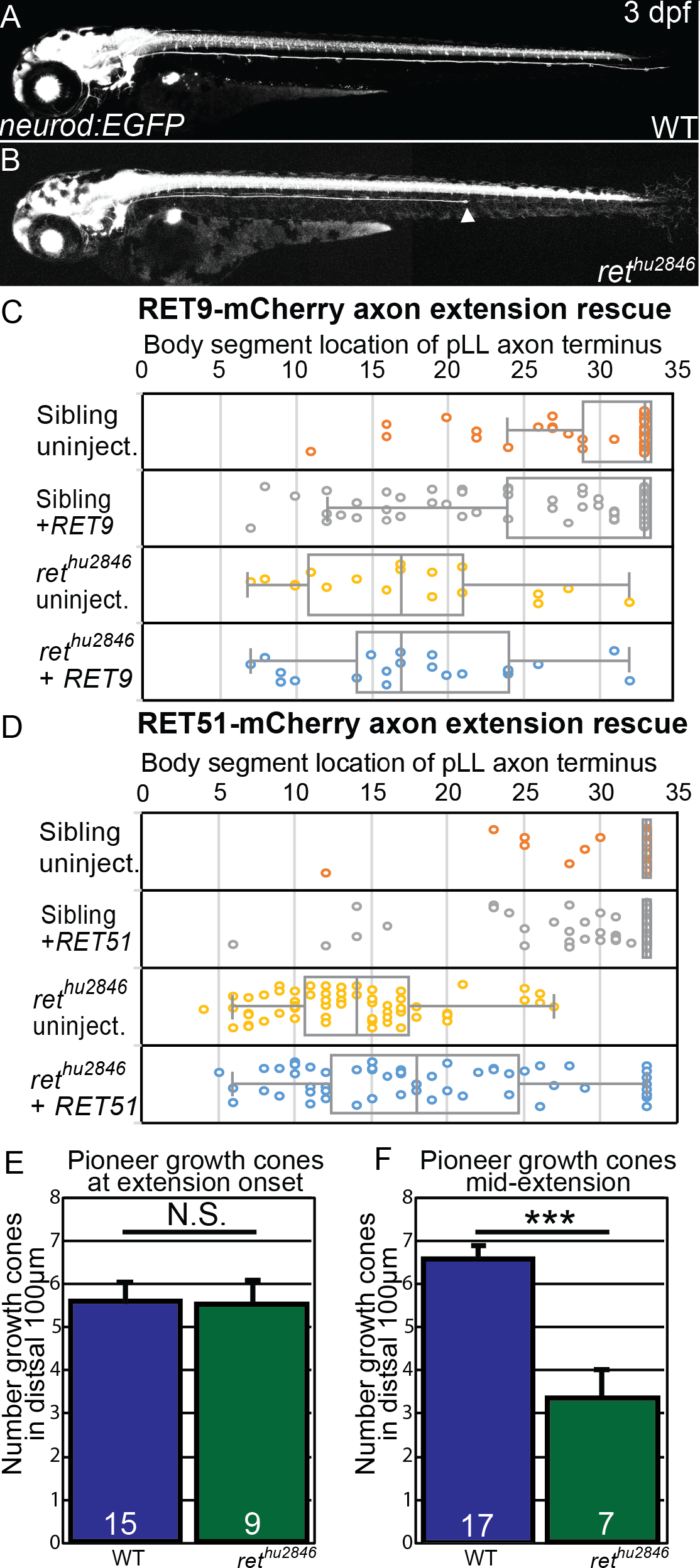
*ret51* is required for pioneer pLLg axon outgrowth. (A,B) Live lateral view of wild-type siblings (A) and *ret^hu2846^* mutants (B) with neurons labeled by the TgBAC(*neurod:EGFP)* transgene. *ret^hu2846^* mutants display truncated pLLg axons (arrowhead). (C,D) Whisker plots comparing mRNA rescue of pLLg axon truncation defects within the same clutches of *ret^hu2846^* mutants with injection of *RET9-mCherry* (C) or *RET51-mCherry* (D).*RET9-mCherry* injection did not significantly ameliorate pLLg axon truncation (sibling uninjected n=50, sibling + *RET9* n=98, *ret^hu2846^* uninjected n=18, *ret^hu2846^ + RET9* n=21); in contrast, *RET51-mCherry* significantly increased mean pLLg axon length as well as cases of bilateral, full-length axon extension rescue (sibling uninjected n=70, sibling + *RET51* n=166, *ret^hu2846^* uninjected n=61, *ret^hu2846^ + RET51* n=87). (E,F) Number of pioneer pLLg axon growth cones in sibling and *ret^hu2846^* mutant embryos at 24 hpf (E) and 30 hpf (F) detected by a-SCG10 immunostaining. *ret^hu2846^* mutants display a significantly lower number of pioneer growth cones at 30 hpf, but not 24 hpf. Error bars represent S.E.M., *** = *p*\*<0.001, N.S. = not significant.

### The Ret51 isoform is required for lateral line axon extension

Because Ret is elevated in pioneer neurons during extension, we asked whether Ret activity is required for pioneer axon outgrowth, and, if so, which Ret isoform is necessary for this process. Using the *neurod:EGFP* transgene to visualize the pLLg axons, we observed that mutants homozygous for the null *ret^hu2846^* allele^31^ (which leads to loss of Ret9 and Ret51 isoforms) had disrupted pLLg axon extension, ultimately producing prematurely truncated axons (Fig. 2B). We observed 97.4% of homozygous *ret^hu2846^* mutants had premature truncation in one or both pLLg axons by 3 dpf (n=89), though we also observed variability in truncation severity between embryo clutches collected from different adult pair matings. However, within a particular clutch, *ret^hu2846^* phenotypes were consistent, as evidenced by the small standard error of the mean of axon truncation in *ret^hu2846^* mutants in subsequent rescue experiments (Fig. 2C,D). Because of pair to pair variability, we compare *ret^hu2846^* mutant embryos to sibling controls within clutches from individual pair matings. To test if a particular Ret isoform was required for pLLg axon outgrowth, we injected *ret^hu2846^* mutant and sibling embryos from individual clutches with *in vitro* transcribed human Ret isoform mRNA tagged with mCherry, either *RET9-mcherry* or *RET51-mcherry*^18^, at the one-cell stage and assayed at 3 dpf for axon length, measuring the shortest pLLg axon bundle in the embryo as a conservative measure of axon extension rescue (Fig. 2C,D). Comparing uninjected *ret^hu2846^* mutant embryos to *RET* isoform mRNA-injected *ret^hu2846^* mutant siblings within clutches we found *RET51*, but not *RET9*, significantly ameliorated the pLLg axon truncation phenotype (*ret^hu2846^* mutant uninjected mean body segment location of axon termination: 17.5±1.7 vs. *ret^hu2846^* mutant *RET9* mRNA injected: 18.1±1.6, *p*=1.00 by two-way ANOVA, see Methods for further details; *ret^hu2846^* mutant uninjected: 13.8±0.7 vs. *ret^hu2846^* mutant *RET51* injected: 18.8±0.9, *p*<0.001). In some cases, *RET51* injection produced full-length, bilateral rescue of axon extension (Fig. 2D, 9 of n=87 embryos). This suggests that, in contrast to other developmental contexts such as enteric nervous system and kidney development where Ret9 is necessary but Ret51 is largely dispensable, Ret51 is the critical isoform required for Ret-mediated pLLg axon outgrowth.

### Ret is not required for sensory pioneer neuron differentiation, but is necessary for proper growth cone morphology

Because pLLg axon extension is incomplete in *ret^hu2846^* mutants, we examined the cellular bases of this axon growth failure. After initial axon outgrowth and premature termination of axon extension in *ret^hu2846^* mutant embryos, we observed no degeneration or retraction of the partially extended pLLg axons from 48 hpf through 120 hpf (Suppl. Fig. 1). To determine if there was a difference in the number of pioneer neurons that extended axons during initial outgrowth, we immunostained *ret^hu2846^* mutants and their wild-type siblings with a-SCG10 antibody at 24 (early outgrowth) or 30 hpf (mid-outgrowth) and imaged distal pLLg axons. SCG10 (STMN2) is enriched in extending sensory pioneer growth cones^32^, serving as a marker of individual pioneer growth cones. Based on SCG10 expression in the distal 100 µm of axons (visualized by the *neurod:EGFP* transgene) the number of pioneer growth cones that initially extended from the pLLg at 24 hpf was not significantly different between *ret^hu2846^* mutants and siblings (Fig. 2E, wildtype: 5.73±0.29 vs. *ret^hu2846^*: 5.56±0.48, *p*=0.99 by Mann-Whitney U test), indicating there is no defect in pioneer neuron specification or initial axon outgrowth in *ret^hu2846^* mutants. However, at 30 hpf, *ret^hu2846^* mutants had significantly fewer growth cones in the distal 100 µm of the growing pioneer axon bundle (Fig. 2F, wildtype: 6.58±0.32 vs. *ret^hu2846^*: 4.00±0.44, *p*<0.001). These observations suggest that pLLg pioneer neurons are specified and begin to extend normally; however, axon extension fails after initial outgrowth.

To explore the effect of Ret loss-of-function on pioneer growth cone morphology and behavior, we assayed axon terminals of extending pioneer axon bundles in *ret^hu2846^* mutants and wild-type siblings (Fig. 3A,B). First, we quantified axonal volume of the distal 75 µm of *neurod:EGFP* transgenic embryos at 30 hpf which typically contains 5-6 pioneer growth cones (Fig. 3C). In comparison to siblings, *ret^hu2846^* mutants had significantly reduced distal axonal volume in extending pioneer axon bundles (Fig. 3C; wildtype: 1477.6±48.9 µm^3^ vs *ret^hu2846^*: 1137.4±136.1, *p*=0.003). To visualize individual pioneer axon and growth cone morphology, *neurod:EGFP* embryos at the one-cell stage were injected with plasmid containing the *neurod5kb* promoter^33^ driving mCherry (*neurod5kb:mCherry)*. Injected plasmid distributes mosaically in zebrafish embryos, allowing identification and live imaging of individual pLLg pioneer growth cones expressing mCherry during extension (30 hpf) (Fig. 3A’,B’). Compared to wild-type siblings, *ret^hu2846^* mutant pioneer growth cones were much less elaborate, had significantly reduced volume (Fig. 3D; wildtype: 291.5±24.2 µm^3^ vs *ret^hu2846^*: 182.6±17.4, *p*<0.001), and significantly fewer filopodia (defined as protrusions ³1 µm) per growth cone (Fig. 3E, wildtype: 9.4±0.7 vs. *ret^hu2846^*: 5.9±0.6, *p*<0.001). However, the length of these filopodia was not significantly different (Fig. 3F, wildtype: 4.55±0.22 µm vs. *ret^hu2846^*: 4.08±0.29, *p*=0.20) between the two groups. Together, these data show that Ret signaling is required for maintaining proper pioneer growth cone morphology and filopodial numbers but not length.

**Figure 3:**
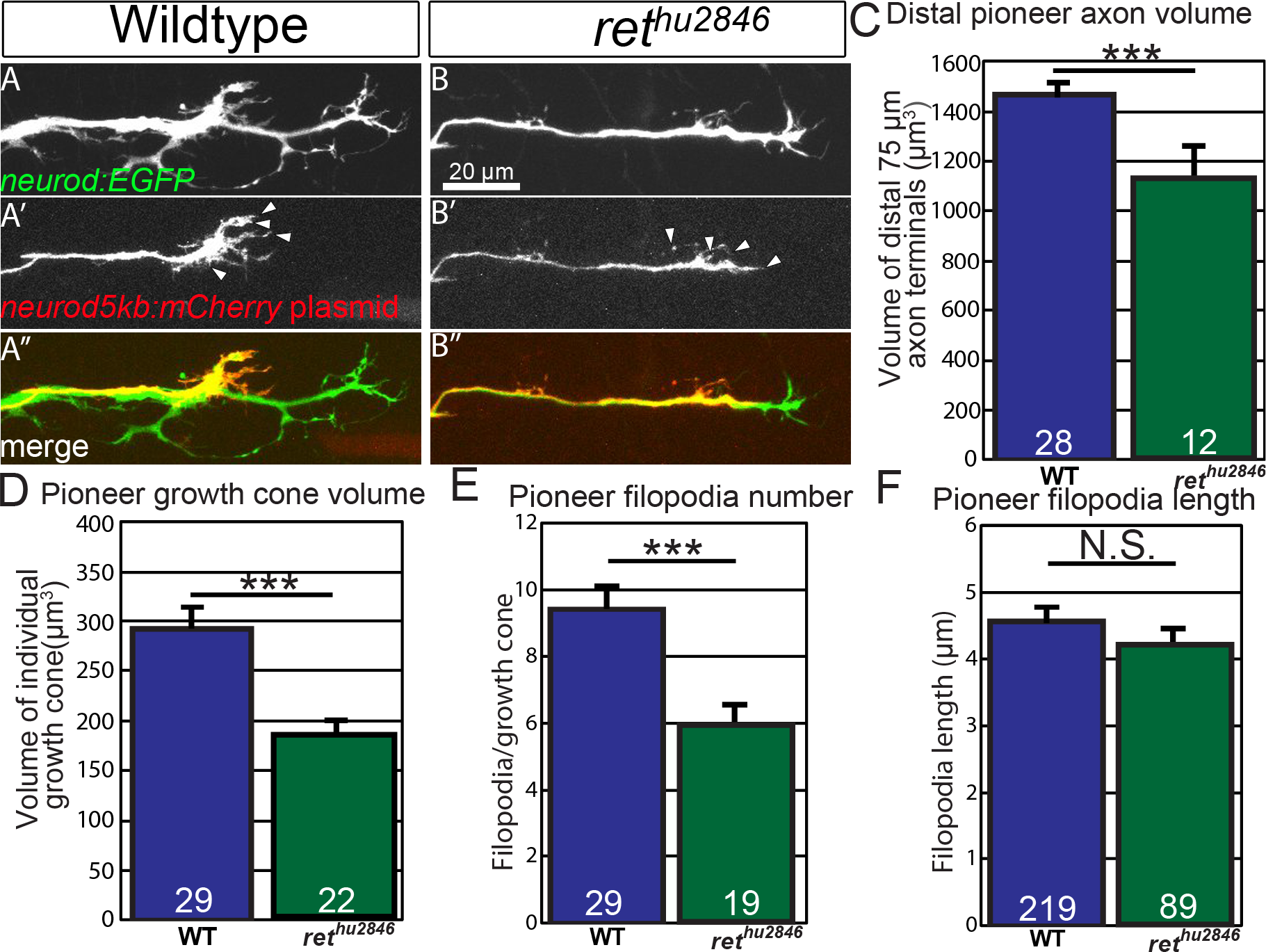
Ret loss-of-function alters pioneer axon growth cone morphology and protrusion. A,B) Lateral view, live image of collective pioneer axon terminals of wild-type siblings (A) and *ret^hu2846^* mutants (B) labeled by TgBAC(*neurod:EGFP*) transgene at 30 hpf. (A’,B’) Individual pioneer axons labeled by mosaic expression of *neurod5kb:mCherry* plasmid (Arrowheads = examples of counted filopodia). (C,D) Quantification of distal collective axon terminal volume (C) or individual pioneer growth cone volume (D). *ret^hu2846^* mutants have significantly reduced collective axonal and individual growth cone volume. (E,F) Quantification of number (E) and length (F, n=filopodia counted) of filopodia (≥1 µm) per individual pioneer axon. Note that *ret^hu2846^* mutants have significantly reduced number of filopodia but not length. Error bars represent S.E.M., *** = *p*<0.001, N.S. = not significant.

### Jip3 loss-of-function phenocopies loss of Ret signaling

Neurotrophin signaling often involves retrograde transport of the activated receptor-ligand complex from the axon terminal to the cell body. However, the specific trafficking mechanisms are not clearly understood. In the pLLg, JNK-interacting protein 3 (Jip3) functions as a specific retrograde cargo adaptor for Dynein-mediated axonal transport during development^34^. Embryos homozygous for the Jip3 null allele, *jip3^nl7^*, display pLLg axon extension defects similar to *ret^hu2846^* mutants (Fig. 4B)^34^. Thus, we investigated whether Jip3 and Ret act in the same pathway during axon outgrowth. We discovered that, similar to *ret^hu284^* mutants, *jip3^nl7^* mutants displayed reduced terminal pioneer axon volume (4D, E, wildtype: 1294.7±51.5 µm^3^ vs *jip3^nl7^*: 1134.7±57.2, *p*=0.024) and fewer filopodia per pioneer growth cone (Fig. 4G, wildtype: 10.3±0.9 vs. *jip3^nl7^*: 7.7±0.4, *p*=0.017) with no significant change in filopodial length (Fig. 4H, wildtype: 3.88±0.38 µm^3^ vs. *jip3^nl7^* mutant: 4.24±0.26, *p*=0.33). Additionally, double homozygous *ret^hu2846^* mutant and *jip3^nl7^* mutants had no significant difference in axon truncation defects compared to single homozygous *ret* mutants (Fig. 4I, body segment location of axon termination in *ret^hu2846^* mutants, +/+: 14.4±1.7, *jip3^nl7^/+*: 13.9±3.0, *jip3^nl7^/jip3^nl7^*: 15.7±3.9, *p*=0.92 by one-way ANOVA), indicating Jip3 and Ret operate in the same genetic pathway. Based on these findings, we investigated whether loss of Jip3 affects localization of Ret protein in the extending axons. *jip3^nl7^* mutants displayed significant accumulation of both total and p905 Ret receptor (referred to as tRet and pRet, respectively; for a-pRet antibody validation, see Suppl. Fig. 3) at the pioneer growth cone during extension compared to wild-type siblings (Fig. 5A-F, for tRet, wildtype fluorescence intensity: 1.97±1.15 vs. *jip3^nl7^*: 45.09±12.44, *p*=0.033; for pRet, wildtype: 7.59±1.60 vs. *jip3^nl7^*: 20.54±3.90, *p*<0.001). These data suggest that a pool of activated Ret receptor fails to be trafficked from the extending growth cone in *jip3^nl7^* mutants.

**Figure 4:**
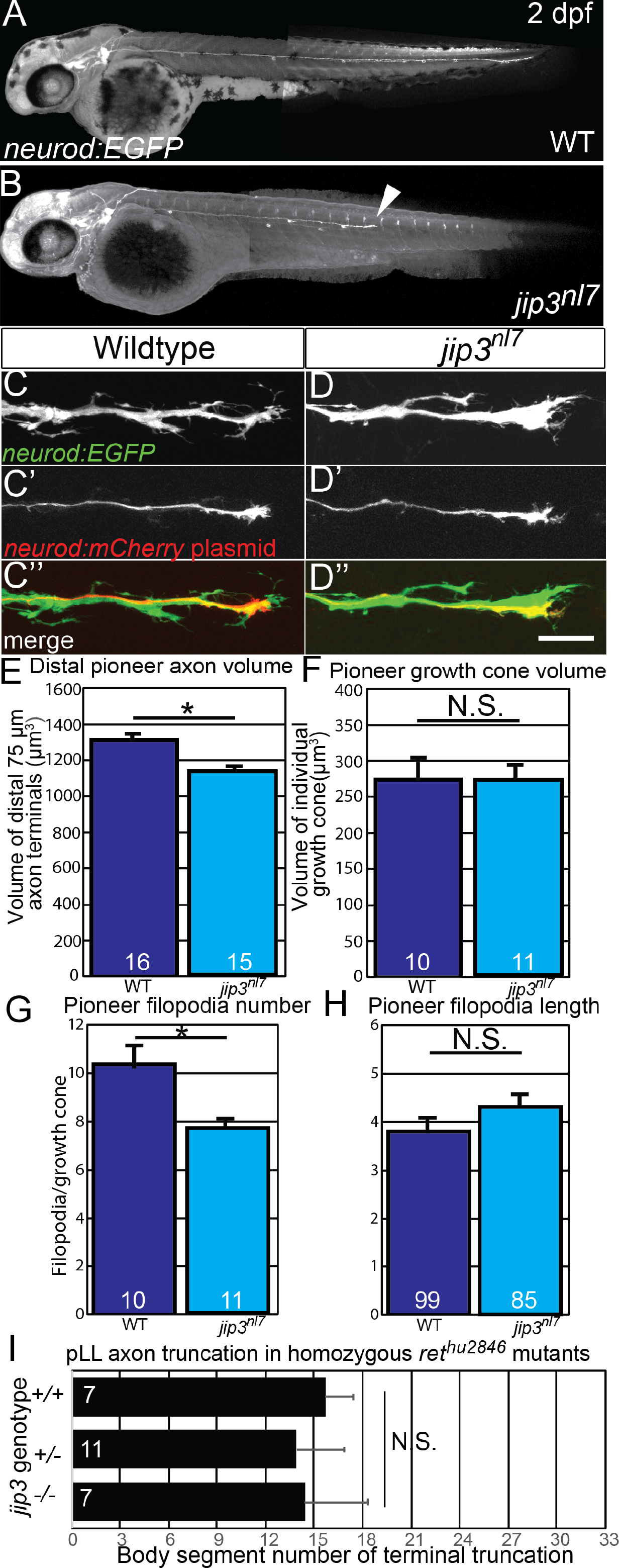
Loss of Jip3 phenocopies *ret* mutant pioneer axon defects. (A,B) Lateral view, live image of wild-type sibling (A) and *jip3^nl7^* mutant (B) containing the TgBAC(*neurod:EGFP*) transgene at 2 dpf. *Jip3^nl7^* mutants display pLLg axon truncation defects (arrowhead). (C,D) Lateral view, live image of collective pioneer axon terminals of wild-type siblings (C) and *jip3^nl7^* mutants (D) labeled by TgBAC(*neurod:EGFP*) transgene at 30 hpf. Scale bar = 20µm (C’,D’) Individual pioneer axons in the same animal labeled by mosaic expression of *neurod5kb:mCherry* plasmid. (E,F) Quantification of distal collective axon terminal volume (E) or individual pioneer growth cone volume (F). *Jip3^nl^* ^7^ mutants have significantly reduced collective axonal volume but not individual growth cone volume. (G,H) Quantification of number (G) and length (H) of filopodia (≥1 µm) per individual pioneer axon. *Jip3^nl7^* mutants have significantly reduced number of filopodia but not length. (I) Quantification of axon truncation defects of sibling, heterozygous, and homozygous *jip3^nl^* ^7^ mutants in homozygous *ret^hu2846^* background shows no significant difference in axon truncation phenotype between genotypes. Error bars represent S.E.M., * = *p*<0.05, N.S. = not significant.

**Figure 5:**
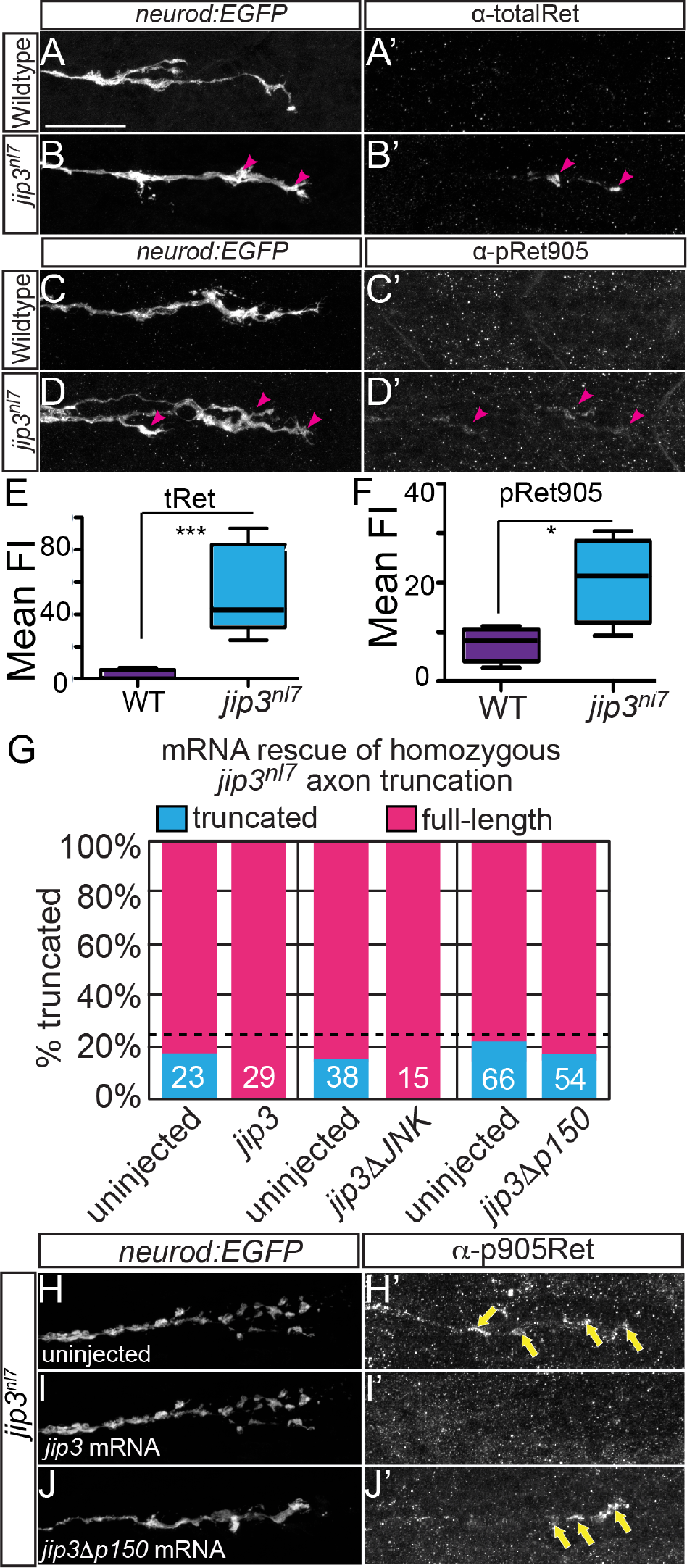
Jip3 is required for clearance of activated Ret from pioneer axon growth cones. (A-D) Lateral view, immunostained pioneer axon growth cones labeled by Tg(*neurod:EGFP*) transgene at 30 hpf. Immunostaining for total Ret receptor (A’,B’, WT n=8, *jip3^nl7^* n=5) and activated pRet905 receptor (C’,D’, WT n=4, *jip3^nl7^* n=4) shows no signal in wild-type sibling pioneer growth cones (A’,C’) but a notable accumulation of total and pRet905 receptor in *jip3^nl7^* mutants (B’,D’). (E,F) Quantification of immunostaining signal in pioneer growth cones displayed in whisker plots. Both total (tRet) and pRet905 signal is significantly higher in *jip3^nl7^* mutant pioneer growth cones. * = *p*<0.05, *** = *p*<0.001. (G) Embryos derived from *jip3^nl7^/+* heterozygous crosses were injected with wild-type *jip3, jip3* lacking the JNK-binding domain (*jip3∆JNK*), or *jip3* lacking the p150-binding (*jip3∆p150*) mRNA. Expression of wild-type and *jip3∆JNK* mRNA completely rescued pLLg axon truncation defects but *jip3∆p150* did not. Dotted line is drawn at 25%, expected fraction of axon truncation in a heterozygous incross. Numbers represent *n* embryos. (H-I) Fixed and immunostained *jip3^nl7^* growth cones at 30 hpf from embryos injected with *jip3* mRNA constructs. Full-length *jip3* mRNA injection (I) rescues pRet accumulated in uninjected pLLg growth cones (H, yellow arrows) but *jip3∆p150* (J) does not.

We next asked whether retrograde transport of Jip3 is necessary for the clearance of Ret from the growth cone and axon extension. To test this, we injected EGFP-tagged mRNAs encoding full-length Jip3 (control), Jip3 lacking the p150 binding domain that is unable to interact with the Dynein motor complex^35^ (*jip3**Δ**p150*) or Jip3 lacking the JNK interacting domain^36^ (*jip3**Δ**JNK*) into 1-cell stage embryos from a heterozygous +/*jip3^nl7^* mutant incross (Fig. 5G). Embryos were fixed at 3 days and immunostained for neurofilaments (3A10) to visualize pLLg axons and the proportion of embryos with axon truncation were scored. Expression of a full-length *jip3* completely rescued axon truncation at 3 dpf (Fig. 5G, uninjected truncation rate: 17.4%, n=23 vs. *jip3*: 0%, n=29). Within a clutch, *jip3**Δ**JNK* also rescued truncation while *jip3**Δ**p150* failed to rescue (uninjected: 15.8%, n=38, *jip3**Δ**JNK*: 0%, n=15, *jip3**Δ**p150*: 22.2%, n=36). Additionally, expression of full-length *jip3* mRNA rescued accumulation of pRet in pioneer growth cones (Fig. 5I) but *jip3**Δ**p150* did not (Fig. 5J). To test if Ret signaling locally regulates JNK activation, we compared immunostaining for pJNK in pioneer growth cones at 30 hpf between *ret^hu28^46* mutants and siblings, but found no significant difference (Suppl. Fig. 4). Altogether, these data indicate that the Jip3-p150 interaction is required for Jip3-dependent axon extension and clearance of pRet from pLLg growth cones, suggesting disrupted Ret retrograde transport may underlie the axon truncation observed in *jip3^nl7^* mutants.

### Jip3 mediates retrograde transport of activated Ret51 receptor

Based on these phenotypic similarities and the accumulation of activated Ret in growth cones of *jip3^nl7^* mutants and the role of Jip3 as a retrograde transport adapter, we asked if Jip3 mediates retrograde transport of Ret. To address this question, we first performed nerve sever experiments^34^ as an indirect readout of Ret axonal transport (Fig. 6A-F). Post nerve sever, anterogradely transported cargoes accumulate in the proximal stump whereas retrogradely transported cargoes accumulate in the distal stump^34^. We hypothesized that if Jip3 specifically mediates retrograde axonal transport of Ret from the growth cone, Ret protein should accumulate at the distal site of nerve sever. Extended pLLg pioneer axons (labled by *neurod:EGFP*) of siblings and *jip3^nl7^* mutants were cut between NM 2 and 3 at 5 dpf, allowed to recover for 3h, fixed, and immunostained for total and p905 Ret. Total Ret was detected at the proximal and distal cut sites, indicating that Ret is bidirectionally trafficked in axons. However, the amount of total Ret was not different between wild-type and *jip3^nl7^* mutant embryos (Fig. 6A-D, proximal fluorescence intensity, wildtype: 32.0±2.5 vs. *jip3^nl7^*: 28.4±3.5, p=0.39; distal FI, wildtype: 26.2±2.3 vs. *jip3^nl7^*: 28.4±2.8, p=0.54), arguing that Jip3 is not required for this process. In wild-type siblings and *jip3^nl7^* mutant embryos, we observed no significant accumulation of pRet at the proximal stump (Fig. 6C,F, wildtype: 9.0±2.0 vs. *jip3^nl7^*: 9.8±3.1, *p*=0.82); however, pRet accumulated significantly at the distal stump of wild-type siblings (Fig. 6C’) compared to *jip3^nl7^* mutants, which had much less pRet accumulation at the distal cut site (Fig. 6D’,F, wildtype: 22.4±2.9 vs. *jip3^nl7^*: 11.4±1.8, *p*=0.004). These data imply that Jip3 is required for retrograde transport of activated Ret.

**Figure 6.**
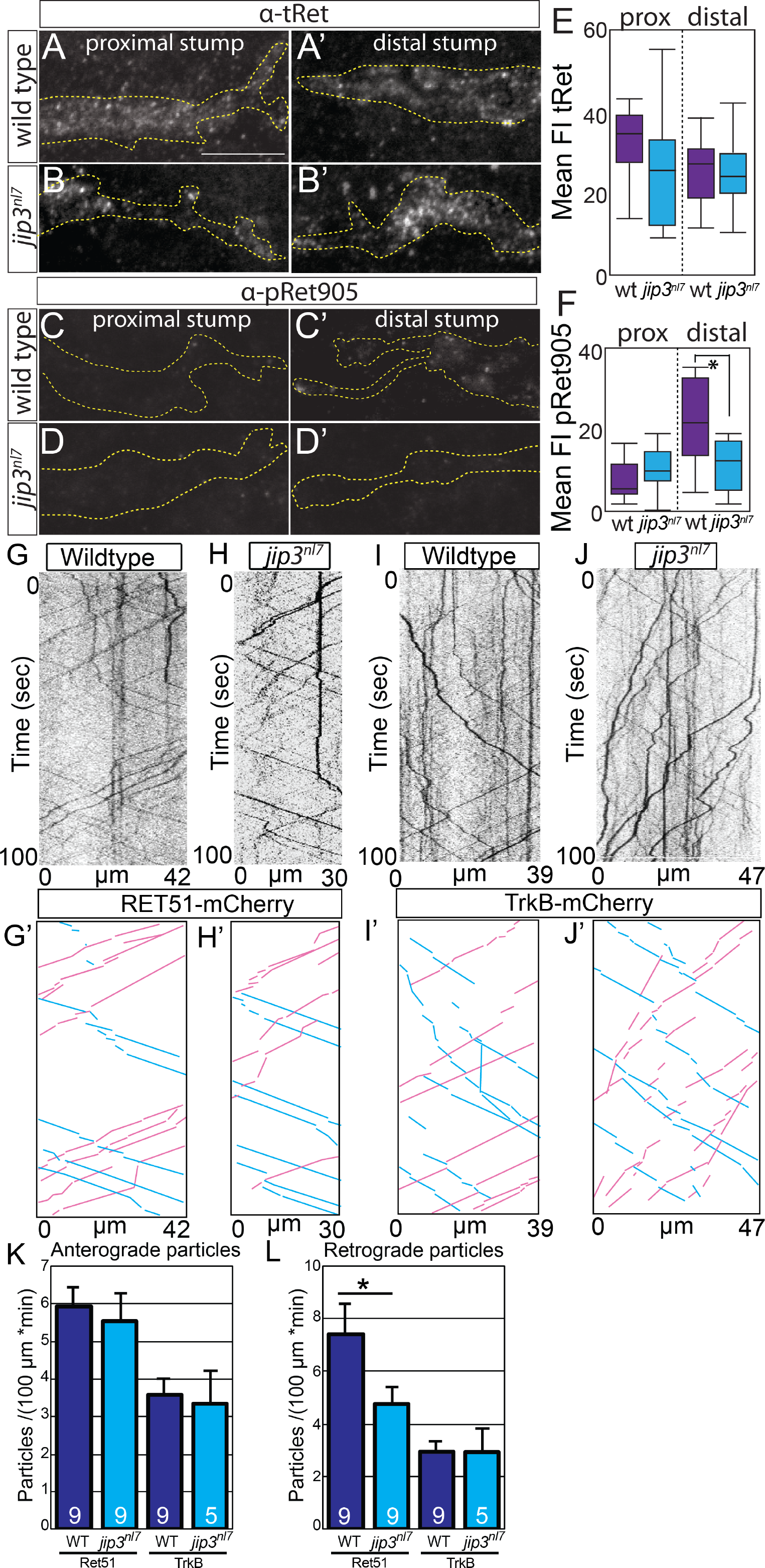
Jip3 mediates retrograde axonal transport of activated Ret51. Lateral view, immunostaining of severed pLL nerves proximal (A-D) and distal (A’-D’) to injury site, 3 hours post-injury; dotted line = nerve outline. (A,B) tRet immunostaining is not notably different between wild-type siblings (A,A’) and *jip3^nl7^* mutants (B,B’) on either side of injury site. (C,D) p905Ret staining is mostly absent in wild-type siblings proximal to injury site (C) and *jip3^nl7^* mutants on both sides of the injury site (D,D’). Stronger p905Ret staining signal is present in wild-type siblings distal to the injury site (C’), indicating presence of retrograde transport of activated Ret receptor in the axon. (E,F) Quantification of tRet (E) and p905Ret (F) staining following axon sever (WT n=12, *jip3^nl7^* n=14) displayed by whisker plot (* = p<0.05). Note p905Ret signal in wild-type siblings is significantly higher than in *jip3^nl7^* mutants, suggesting a failure of retrograde transport of pRet receptor in mutants. (G-J) Kymographs of trafficking in individual pLLg axons of Ret51-mCherry (G,H) and TrkB-mCherry fusions (I,J) in wild-type siblings and *jip3^nl7^* mutants. (G’-J’) Kymograph traces of scored anterograde (blue) and retrograde (magenta) transported particles. (K) Quantification of normalized anterograde particle counts from kymograph analysis (wildtype Ret51=5.93±0.52 vs. *jip3^nl7^* Ret51=5.53±0.76, *p*=0.86 by Mann-Whitney U test; wildtype TrkB=3.58±0.42 vs. *jip3^nl7^* TrkB=3.36±0.86, *p*=0.82). (L) Quantification of normalized retrograde particle counts from kymograph analysis (wildtype Ret51=7.33±1.17 vs. *jip3^nl7^* Ret51=4.73±0.53, *p*=0.36; wildtype TrkB=3.07±0.31 vs. *jip3^nl7^* TrkB=3.06±0.75, *p*=1.00). *jip3^nl7^* mutants show a significant decrease in the number of retrogradely transported Ret51-mCherry particles but no change in TrkB-mCherry particle counts or anterograde Ret51-mCherry particles. * = *p*<0.05. Error bars represent S.E.M.

Next, we tested directly if Jip3 is required for axonal transport of Ret receptor by observing trafficking of Ret fusion protein in extending pLLg axons. To achieve this, we injected plasmid driving mCherry-tagged RET9 or RET51 under the *neurod5kb* promoter (*neurod5kb:RET9/51-mCherry)* at the one-cell stage and imaged transport of mCherry-positive particles in extending pLLg axons at 30 hpf^37^. Injected plasmid distributes mosaically in the developing embryo, thus, we can select embryos expressing fusion constructs in 1-3 pLLg neurons to assay Ret trafficking in individual axons during extension. We observed anterograde and retrograde trafficking of mCherry-positive particles in *neurod5kb:RET51-mCherry* injected embryos, but did not observe any trafficking of discrete mCherry-positive particles with injection of *neurod5kb:RET9-mCherry* (data not shown). To test whether Jip3 regulates Ret51 retrograde axonal transport in pioneer neurons, we assayed *RET51-mCherry* transport in *jip3^nl7^* embryos and wild-type siblings. For our experiments, we used expression of the *neurod:EGFP* transgene to identify the most distal pLLg pioneer neurons also expressing RET51-mCherry (Fig. 6G,H). Based on kymograph analysis, we found significantly fewer retrogradely trafficked Ret51-mCherry particles in *jip3^nl7^* mutant pLLg pioneer neurons compared to wild-type siblings, suggesting a disruption in retrograde transport of Ret51 (Fig. 6G’,H’,K). To examine whether transport of other neurotrophin receptors expressed in the pLLg, such as TrkB^38^, was disrupted, we analyzed trafficking of TrkB-mCherry fusion from embryos injected with *neurod5kb:trkB-mCherry* plasmid (Fig. 6I,J). Notably, we saw no significant change in trafficking of TrkB-mCherry particles (Fig. 6K,L). These data, taken together with pRet accumulation in the distal stump of severed *jip3^nl7^* mutant pLLg axons, strongly argue that Jip3 is required for transport of activated Ret51, but not TrkB, in pLLg axons.

Finally, if Jip3 acts as a specific cargo adaptor for retrograde pRet51 transport, we expect that Jip3 and Ret51 would be co-transported in extending pLLg axons. To address this, we co-injected *Tg(BAC)RET51-EGFP* and *neurod5kb:jip3-mCherry^34^* plasmid at one-cell stage and imaged pLLg pioneer axons expressing both fusions at 30 hpf. Co-localization of Jip3-mCherry and RET51-EGFP was observed in a subset of retrogradely trafficked particles but no cases of co-localization were observed for anterogradely transported particles (Fig.7, Suppl. Mov. 1). Taken together, these data suggest Jip3 preferentially mediates retrograde transport of activated pRet51.

**Figure 7:**
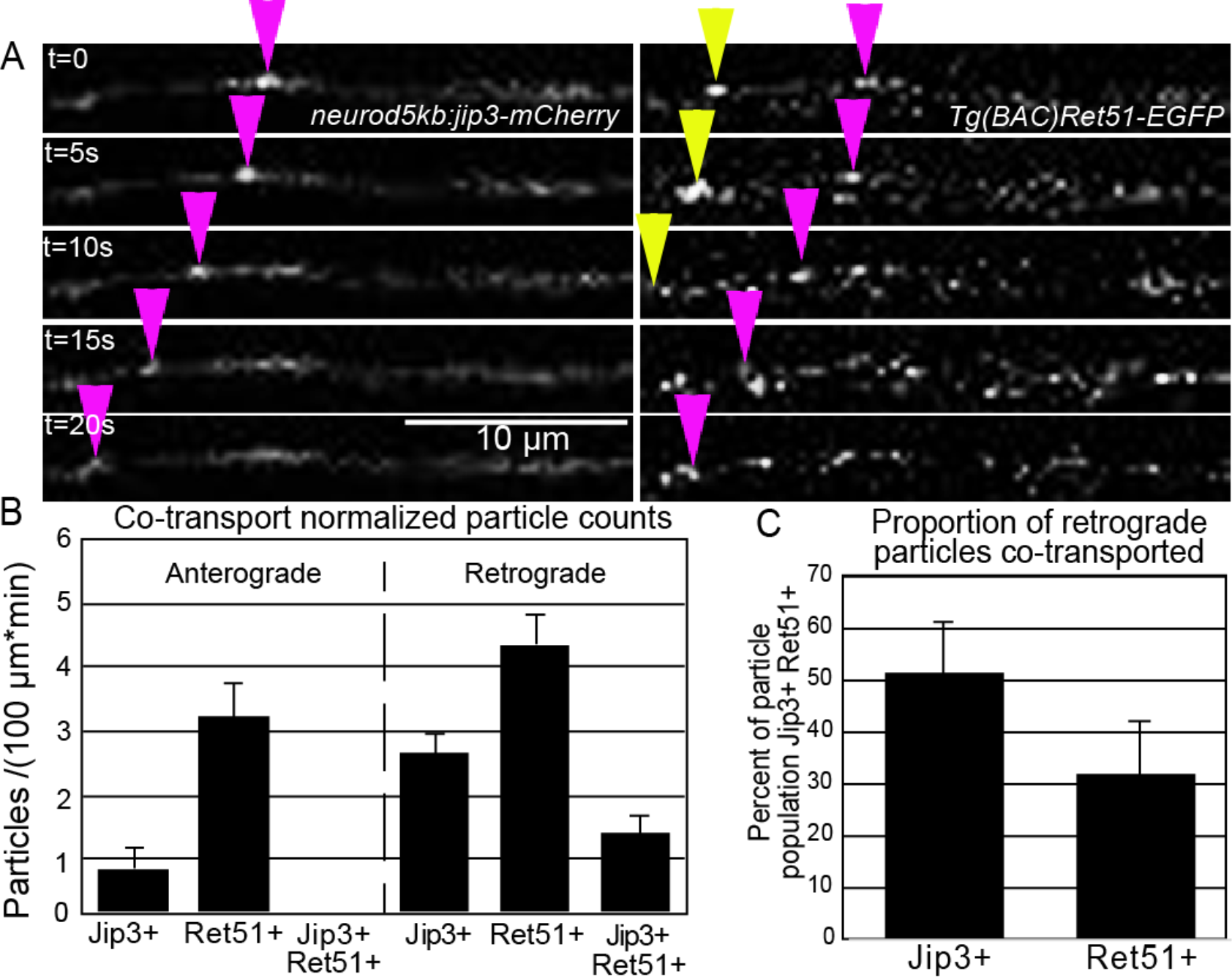
Ret51 and Jip3 co-transport in extending pLLg axons *in vivo*. (A) Still images from Supplementary Movie 1 of pLLg axonal transport in embryos co-injected with *neurod5kb:jip3-mCherry* and Tg(BAC)*Ret51-egfp* plasmids. Retrograde transport of mCherry+ eGFP+ particles can be seen (magenta arrowhead) as well as solely GFP+ particles (yellow arrowhead). (B) Quantification of Jip3-mCherry and Ret51-eGFP normalized particle counts (n= 4 embryos). Anterograde Ret51+ Jip3+ particles were not observed, however retrograde Ret51+ Jip3+ particles were seen. (C) Proportion of Jip3+ or Ret51+ particle populations that are also Jip3+ and Ret51+. Approximately half of Jip3+ retrograde particles are also Ret51+. Error bars represent S.E.M.

## Discussion

The important developmental role of pioneer neurons and their unique axon outgrowth and pathfinding capabilities have led to their study in the CNS and PNS of vertebrates and invertebrates. Despite the wide variety of molecular and genetic tools available for neurobiological research, pioneer neurons are still largely identified at an anatomical level using differences in growth cone morphology, timing of axon outgrowth, or the stereotyped anatomical location of cell bodies in the case of well-mapped nervous systems, such as *Drosophila*. The lack of a strong, specific molecular or transcriptional marker precludes many avenues of study to elucidate the fundamental neurobiological mechanisms that promote the axon growth and pathfinding behaviors of pioneer neurons. Our work identifies Ret51 as a molecular marker of pioneer sensory neurons and a regulator of pioneer axon outgrowth. We observed high expression of *ret51* mRNA specifically in a region of the pLLg neurons known to hold the pioneer neuron cell bodies^29^ compared to the broad expression of *ret9* throughout the pLLg. Through retrograde dye labeling of pioneer neuron cell bodies, we found an enrichment of Ret receptor protein in pLLg pioneer neuron cell bodies, corresponding to the region of *ret51* mRNA expression. *ret^hu2846^* mutants do not have defects in pLLg pioneer neuron specification or survival but do have a failure of continued growth of pioneer axons. This pioneer axon growth was rescued in *ret^hu2846^* mutants by exogenous expression of *RET51* mRNA but not *RET9*, indicating that Ret51 is specifically required for pioneer axon outgrowth. From these data, we conclude Ret51 is a specific marker of pioneer neurons in the pLLg.

### A specific role for Ret51 signaling in pioneer axon outgrowth

Ret receptor isoforms have different roles and requirements during development. Ret9 is the critical isoform in the vast majority of cases, most notably in enteric nervous system and kidney development. RET51 is expressed in a specific subset of RET9-expressing sensory neurons in the mouse olfactory system during embryonic development^39^, but the functional consequence of this is unknown. Neuronal studies examining differences in Ret9 and Ret51 subcellular localization and degradation have focused on their roles in neuronal survival^16^. Our work establishes a novel, specific role for Ret51 in pioneer axon extension unrelated to neuronal differentiation or survival and describes *in vivo* observation of axonal Ret isoform trafficking. Live imaging of RET51/Jip3 retrograde co-transport as well as the disruption of RET51 retrograde axonal transport and pRet localization followed by loss of Jip3, suggests Ret51 retrograde transport may be required for this signaling role. In other cell types, Ret51 is much more rapidly internalized from the plasma membrane and differentially ubiquitylated compared to Ret9^18,40^. In mouse sympathetic neuron culture, Ret51 is more rapidly degraded in axon terminals compared to Ret9 following GDNF treatment^16^. Interestingly, in DRG sensory neuron cultures, Ret9 was much more highly expressed than Ret51, however, Ret51 was substantially more resistant to degradation and activated Ret51 persisted much longer than in sympathetic neurons, potentially allowing long range axonal transport of Ret51 in sensory neurons. Consistent with these results, loss of Jip3 leads to significant accumulation of pRet receptor in pioneer sensory growth cones, suggesting there is not normally a substantial level of degradation of activated Ret receptor locally. Though we observe broad expression of *ret9* throughout pLLg sensory neurons, we were unable to observe axonal trafficking of discrete RET9 particles and *RET9* mRNA failed to rescue mutant axon truncation, suggesting retrograde transport of Ret9 is not required for axon growth. It may be that Ret9 has a role in local signaling at the dendrites or cell body that is unrelated to axon outgrowth. After failure of pioneer axon extension in *ret^hu2846^* mutants, we do not observe axonal degeneration or cell death, indicating that, in contrast to the typical role of Ret in cells with stable terminals at their target, Ret9 or Ret51 are not required for neuronal survival at least through day 5 of development.

### Jip3 is an adapter for transport of activated Ret

While there is evidence that long distance GDNF signaling is required for motor axon pathfinding^25^, to date there has been no *in vivo* observation of retrograde axonal trafficking of Ret receptor or description of the mechanisms that would underlie this transport in neurons^26^. GDNF expression from the pLLp, which guides extending pLLg pioneer axons, is required for proper pLLg axon extension^28^, but the mechanisms by which GDNF/Ret mediate this axon extension are unexplored. We found that loss-of-function of a specific axonal retrograde cargo adapter, Jip3, phenocopied pLLg axon truncation and many of the growth cone defects we observed in *ret^hu2846^* mutants. Jip3 can regulate axonal trafficking of another neurotrophin receptor, TrkB^41^, and acts as a retrograde cargo adapter for multiple cargoes in extended pLLg axons^34^, but has not been implicated in Ret receptor transport or signaling. We observed no difference in axon truncation defects in *jip3^nl7^* and *ret^hu2846^* null double mutants compared to *ret^hu2846^* mutants, indicating that Ret and Jip3 act in the same pathway to promote axon growth. Based on these observations, it is possible Jip3 mediates local Ret signaling in pioneer growth cones, since it also can function as a scaffold that regulates local JNK phosphorylation. However, while Jip3 interaction with JNK has been previously implicated in axon extension^42^, it is not necessary for pLL axon extension^34^ and we observe no change in pJNK localization in pioneer growth cones in *ret^hu2846^* mutants. Consistent with this, we found that mRNA injection of full-length Jip3 and Jip3 lacking the JNK-binding domain in *jip3^nl7^* mutants completely rescued axon truncation and accumulation of pRet in pioneer growth cones, however, Jip3 lacking the p150-binding domain (required for association with the Dynein motor complex) is unable to rescue either phenotype. This suggests Jip3 requires the ability to be retrogradely transported in order to mediate Ret signaling in this context. We observed a significant reduction in the number of retrogradely trafficked Ret51 particles in *jip3^nl7^* mutants but found normal transport of TrkB in *jip3^nl7^*mutants. Finally, we observed live retrograde co-transport of Jip3 and Ret51 tagged particles in extending pLLg axons, but no co-transport of anterogradely trafficked particles. We conclude from these data that Jip3 specifically mediates Ret51 retrograde transport. Notably, we did not observe complete loss of retrograde transport of Ret51 in *jip3^nl7^* mutants or observe complete co-localization of Jip3/Ret51 retrograde transport. This is consistent with our assumption that Jip3 is required for retrograde transport of activated Ret51, which presumably constitutes only a subset of Ret51 fusion visualized in our live trafficking experiments.

Several lines of evidence indicated Jip3 specifically mediates retrograde axonal transport of activated Ret receptor. Immunostaining of extending pioneer pLLg axons for total Ret (tRet) and phospho-905Ret (pRet) revealed a significant accumulation of tRet and pRet in distal tips of pioneer growth cones in *Jip3^nl7^* mutants. Additionally, this accumulation of pRet in *Jip3^n^l7* mutants could be rescued by full-length *jip3* mRNA injection, but *jip3* mRNA lacking the p150-binding domain failed to do so. Our nerve sever experiments found pRet normally accumulated at the distal stump site of wild-type embryos, indicating retrograde transport of activated receptor from axon terminals, but there was significant reduction of pRet signal at the distal stump in *jip3^nl7^* mutants. Taken together, these results indicate Jip3 mediates retrograde transport of phosphorylated Ret51 in pioneer axons and is required for axon outgrowth and proper formation of growth cone filopodia.

### Ret signaling regulates pioneer growth cone morphology and dynamics

Regulated growth cone dynamics, including formation and disassembly of filopodia and lamellipodia, is required for growth cone advancement and axon outgrowth. With Ret loss-of-function, we observe a failure of pioneer axon outgrowth as well as a significant reduction in pioneer growth cone volume. The reduced growth cone volume and more compact morphology of *ret* mutant growth cones which lacks the typical spread out “footprint” of pioneer growth cones could be caused by defects in formation of lamellipodia, which would ultimately prevent proper growth cone migration and steering. Loss of Ret also led to fewer filopodia per pioneer growth cone but did not change mean length of filopodia. This suggests a defect in initiation of filopodial formation but not maintenance or continued growth. Failure to dynamically form filopodia could disrupt proper interaction with the extracellular environment, particularly the pLLp which is producing GDNF and BDNF^38^, necessary for guiding and promoting axon growth. One explanation for defects in axon outgrowth, then, may be that loss of filopodia leads to less interaction with the pLLp and thus a reduction of trophic/permissive cues sensed from the pLLp, further reducing the formation of growth cone filopodia and producing a positive feedback cycle that ends in failed growth cone advancement. Another non-mutually exclusive possibility is that disruption of dynamic filopodial formation simply prevents proper growth cone motility^43^, leading to failed outgrowth. We conclude, based on the changes we observe in pioneer growth cone morphology that one cellular mechanism underlying failure of axon growth in *ret* mutants is defective dynamic actin cytoskeletal arrangement which prevents proper growth cone advancement and pathfinding.

Jip3 loss-of-function produces many of the same cellular growth cone phenotypes as Ret loss-of-function. Notably, individual *jip3^nl7^* mutant pioneer growth cones were not significantly smaller than wild-type, unlike *ret^hu2846^* mutants, which were reduced in volume. This may be explained by terminal pLLg axon swelling that occurs in *jip3^nl7^* mutants due to failures in retrograde transport of specific cargoes^34^. Such swellings could counteract the reduction of pioneer growth cone volume normally seen with Ret loss-of-function. Alternatively, regulation of this aspect of pioneer growth cone morphology by Ret may either be a local event or otherwise not mediated by Jip3. It is also possible that the accumulation of pRet in *jip3^nl7^* mutant growth cones leads to high levels of local Ret signaling that may have an additional local impact on growth cone morphology.

While it is clear that Jip3 retrogradely transports pRet, we propose that long range retrograde Ret signaling mediated by Jip3 is required for pioneer axon outgrowth and growth cone dynamics, as opposed to local effects, for several reasons. First, Jip3 and Ret loss-of-function axon truncation and growth cone phenotypes are the same, with the exception of individual growth cone volume, which could be explained by swelling of axon terminals in *jip3^nl7^* mutants. If it was the case that a local increase in Ret signaling with loss of Jip3 is disruptive to axon outgrowth, we would perhaps expect a difference in axon truncation or filopodial phenotypes compared to *ret^hu2846^* mutants, which we do not observe. Second, there is no difference in the severity of axon truncation in *jip3^nl7^/ret^hu2846^* double mutants compared to individual homozygous mutants, suggesting loss of a Ret retrograde cargo adapter is functionally equivalent to loss of Ret signaling in this case. Third, we observe no local changes in pJNK levels in pioneer growth cones with loss of Ret signaling (Suppl. Fig. 4), indicating that Ret does not have a local effect on at least one major pathway that regulates axon growth and growth cone dynamics. Finally, in the case of other neurotrophins, particularly NGF, retrograde axonal transport is coupled with a role in long range retrograde signaling and not simply a clearance of activated/internalized receptor from the growth cone, which can be regulated by local proteasomal degradation (including in the case of Ret)^16,26,44x^. Based on the established paradigm of retrograde neurotrophin signaling it would be unlikely, then, that activated Ret51 is specifically retrogradely transported along axons without a functional downstream consequence of this transport. Future studies will focus on examining possible effects of high levels of local Ret51 signaling and identifying potential transcriptional targets downstream of Ret retrograde transport and how these regulate specific changes in pioneer growth cone morphology and behavior.

In summary, our *in vivo* data demonstrate a novel role for Ret51 in pioneer axon extension and suggest the requirement of long-range, retrograde Ret signaling in this process. Additionally, we establish a system to examine the role of neurotrophin signaling in pioneer axon growth without a confounding role in neuronal survival. We identify that Jip3 mediates Ret51 retrograde transport, providing a useful tool to dissect local versus long-range signaling contributions of Ret receptor. Combined with our observation that Ret51 is specifically and highly expressed in pLLg pioneer sensory neurons, we submit that Ret51 functions as a molecular and transcriptional marker of these pioneers. This finding opens up several new technical avenues to examine the mechanisms underlying pioneer neuron development and pathfinding. In particular, this finding can be leveraged with transcriptomic techniques, such as single cell RNA-seq, to study the transcriptional mechanisms that specify pioneer neurons and promote their enhanced axon growth and pathfinding capabilities. Better understanding of the fundamental processes that regulate the unique axonal growth and pathfinding capabilities of pioneer neurons can provide potential therapeutic points of intervention to promote axonal regeneration in response to injury or disease.

## Materials and Methods

### Zebrafish husbandry

Adult zebrafish were maintained at 28.5°C as previously described and embryos were derived from natural matings or *in vitro* fertilization^45^, raised in embryo media, and developmentally staged^46^. Strains utilized were AB, TgBAC(*neurod:EGFP)^nl1^ 30*, *ret^hu2846^ 31*, and *jip^nl7^* ^34^.

### Plasmids and tagged constructs

*Neurod5kb:jip3-mCherry* was previously described^*34*^. *Neurod5kb:mCherry* plasmid was generated by recombining the *neurod5kb* promoter^33^ p5e entry vector into the pDEST394 vector in the Gateway system by previously described methods^47^. Human RET9-mCherry and RET51-mCherry constructs in the EGFP-N1 vector were obtained from the Mulligan lab^18^ and digested with HindIII, NotI, and SphI to release the whole tagged construct and digest the EGFP-N1 vector backbone. Released RET fusion construct cassettes were ligated into HindIII/NotI-digested Gateway pME-MCS vector and subsequently recombined with the *neurod5kb* or *cmv/sp6* promoter p5e entry vectors to yield *neurod5kb:RET9/51-mCherry* or *cmv/sp6:RET9/51-mCherry* in the pDEST394 destination vector^47^. Jip3 deletion constructs were derived from pME-Jip3^34^ using the Quickchange II kit (Agilent) with the following primers: **Δ**p150forward (GGGAAAGAAGTGGAAAATGAGGAGCTGGAA TCGGTA), **Δ**p150reverse (TTTACCGATTCCAGCTCCTCATTTTCCACTTCTTTC), **Δ**JNKforward (GAGGAAAAGTAAAACAGGTGGAGATGGCATGGAGGA), **Δ**pJNKreverse (CCATGCCATCTCCACCTGTTTTACTTTTCCTCCAGT). mRNA was synthesized from Jip3 constructs or *cmv/sp6:RET9/51-mCherry* using mMessage Machine (Life Technologies) and microinjected at 500 pg/embryo for Jip3 constructs and 50 pg/embryo for RET9/51-mCherry constructs.

### Generation of the Ret9- and Ret51-eGFP BAC fusions

We modified a Ret-containing bacterial artificial chromosome (BAC) clone by Escherichia coli-based homologous recombination^48^. BAC clone DKEY-192P21 contains 49.89 kb of sequence upstream and 69.78 kb of sequence downstream of *ret* (http://www.sanger.ac.uk/Projects/D_rerio/mapping.shtml). After recombination, the modified BAC clone contained an *eGFP* gene positioned in frame at the 3’ end of exon 19 to generate Ret9 fusion or exon 20 to generate Ret51 fusion, following SV40 polyadenylation signal and kanamycin resistance gene. The kanamycin resistance gene was flanked by FLP sites and was removed by FLP-mediated recombination^48^. The accuracy of EGFP integration and FLP recombination were evaluated by PCR, sequencing, and by transient expression assays.

### In situ hybridization and wholemount immunostaining

RNA *in situ* hybridization was performed as described^49^. Digoxygenin-labeled antisense RNA probes were generated for *ret9* (ensembl: ENSDART00000139237.3) and *ret51* (Refseq: NM_181662.2) by amplifying the alternatively spliced 3’ exons and 3’ UTR of each transcript from cDNA and cloning into pCR4 TOPO (Thermofisher) with the following primers: ret51F (GTCATACCCGAACTGGCCAGA), ret51R (ACTATCGATTGTGTCCACGAT), ret9F (TCCATAGGAAGAGCTGTGATTG), ret9R (GTAAAGACAGGGTTAGTGGCA).

Whole mount immunohistochemistry was performed following established protocols^50^ with the following exception: embryos stained with anti-Ret were fixed in Shandon Glyo-Fixx (Thermo Scientific) for one hour at room temperature. The following antibodies were used: anti-GFP (1:1000; Invitrogen #A11122), anti-SCG10 (1:100; ProteinTech, #10586-AP) anti-p905Ret (1:50; Thermo Pierce, #PA517761), anti-Ret (1:100; Santa Cruz, #sc-167), anti-neurofilament (1:500; DSHB, 3A10) anti-pJNK (1 ∶100; Cell Signaling #9251S). tRet antibody was validated by verifying lack of immunofluorescence signal in pLLg in *ret^hu2864^* mutant embryos compared to wild-type siblings (Suppl. Fig. 2). pRet antibody was validated by injecting a plasmid construct expressing RET51-mCherry fusion (*neurod5kb:RET51-mCherry*), identifying embryos with 2-3 pLLg neurons expressing the construct, fixing at 30 hpf, and immunostaining. Co-localization of Ret51-mCherry and pRet was found in extending growth cones, but pRet was largely absent in the ganglion or cell bodies expressing Ret51-mCherry (Suppl. Fig. 3), consistent with expectation that only a subpopulation of Ret51 would be phosphorylated and that subpopulation would be enriched in extending growth cones near the source of GDNF ligand. All fluorescently labeled embryos were imaged using a FV1000 laser scanning confocal system (Olympus). Brightfield images were collected using a Zeiss Imager Z1 system. Images were processed using ImageJ software^51^. Brightness and contrast were adjusted in Adobe Photoshop and figures were compiled in Adobe Illustrator.

### Quantification of immunofluorescence

For analysis of pRet and tRet intensity in axon terminals and after nerve injury, individuals were immunolabeled as described above. For consistency of labeling, compared larvae were processed in the same batch. Confocal Z-stacks (0.5 µm between planes) were taken of the area of interest using a 40X/NA=1.3 oil objective with identical settings. Images were analyzed using ImageJ^51^. For fluorescence intensity measurements of pRet or tRet in wildtype and mutant growth cones, summed projections of the regions of interest were generated only through regions that contained the *neurod:EGFP* signal and converted to 8 bit in ImageJ as previously described^34^. Briefly, in the pLL nerve injury analysis, a 30 µm, *neurod:EGFP*-positive region encompassing the proximal or distal edge of the severed axon was selected and summed projections through only this segment were compiled for analysis. Prior to statistical comparison, the mean background fluorescent intensity, measured in a region adjacent to the injury site, was subtracted from the values generated.

### Growth cone imaging and analysis

For imaging, embryos were mounted in 1.5% low melting point agarose on a glass coverslip, submerged in embryo media containing 0.02% tricaine (MS-222; Sigma) and imaged at 24 or 30 hpf using a 60X/NA = 1.2 water objective on an upright Fluoview1000 confocal microscope (Olympus). The distal 75-125 µm of pLLg axon terminals, indicated by expression of *neurod:EGFP* transgene, were imaged with stacks of sufficient depth to capture all portions of the terminal axons. For individual growth cone imaging, embryos were injected with *neurod5kb:mCherry* plasmid, embryos expressing mCherry in 1-2 pLLg neurons with axons extending at or near the axon terminals (marked by *neurod:EGFP* expression) were selected and imaged as described using 488 and 568 nm excitation channels. Growth cone and collective axon terminal volume was quantified using Imaris software (Bitplane) as follows. We used EGFP signal to generate axonal surfaces based on full stacks and then calculated the volume encapsulated by the surface in a region of interest containing an individual growth cone or the distal 75 µm of collective axon terminals. Filopodia were visualized by creating Z-projections of image stacks of individual growth cones in ImageJ and filopodia were traced using the segmented line tool, measured for length, and filopodia ≥ 1µm number and length were scored.

### Retrograde axon labeling experiment

40 hpf *neurod:EGFP* embryos were anesthetized and mounted in 1.2% low melt agarose. Tails were severed with rhodamine dextran-soaked scissors just rostral to pLLp growth cones. Embryos were freed from agarose, allowed to develop for 3 hours, then re-mounted and imaged using an FV1000 confocal microscope. Embryos were subsequently fixed and stained to detect Ret immunoreactivity.

### Axon transport analysis

Zygotes were injected with plasmid DNA encoding fluorescently tagged cargos of interest with expression driven by the *5kbneurod* promoter^33^ and imaged as described above. At 30 hpf, embryos were sorted under epifluorescence to identify individuals with tagged cargo expression in a few cells of the pLL ganglion. For each embryo, a region of interest (30–150 µm) was selected in the pLL nerve in which a long stretch of axon was observable in a single plane. Scans were taken at 3 frames per second for 500-1000 frames. Embryos were subsequently released from agarose and processed for genotyping. For co-transport, embryos expressing both constructs in a single cell were selected and imaged as described above using sequential imaging of the 488 and 568 nm excitation channels. 500 frames were collected at 3 frames per second.

Transport parameters were analyzed using kymograph analysis in the MetaMorph software package (Molecular Devices, Inc.) as previously described^37^. Kymographs containing 10 or more traces were analyzed and each were averaged within individual embryos for statistical analysis. The number of particles moving in each direction was estimated based on traces on the kymographs and then normalized to length of axonal segment and total imaging time.

### Axotomy and image acquisition

Five day old zebrafish larvae (*neurod:EGFP* carriers) were anesthetized in 0.02% tricaine and embedded in 3% methylcellulose on a slide. Pulled thick-walled glass capillaries were used to sever the nerve between NMs 2 and 3. Slides were immersed in Ringer’s solution (116 mM NaCl, 2.9 mM KCl, 1.8 mM CaCl_2_, 5 mM HEPES pH = 7.2, 1% Pen/Strep) and incubated at 28.5°C for 3 hours. Larvae were then collected and immunolabeled for pRet or tRet and EGFP.

### Statistical analysis

Data were analyzed with the SPSS software package (IBM). Data suitable for parametric analysis were analyzed using ANOVA with post-hoc Tukey test. Data not suitable for parametric analysis were analyzed using Mann-Whitney *U* test (Wilcoxon ranked sum test). Axon truncation length measuring within groups over time (Suppl. Fig. 1) was analyzed by repeated measures ANOVA with post-hoc Bonferroni test for pairwise comparison. Rescue of Axon truncation length in *ret^hu2846^* mutants by *RET9* or *RET51* mRNA injection was analyzed by two-way ANOVA with post hoc Tukey test for all experimental conditions within a clutch: uninjected wild-type siblings, RET isoform-injected wild-type siblings, uninjected *ret^hu2846^* mutants, and RET isoform-injected *ret^hu2846^* mutants.

## Ethics statement

All animal work was approved by and conducted according to guidelines of the Oregon Health & Science University IACUC.

## Acknowledgements

We thank Dr. Lois Mulligan for her gift of *RET9-mCherry* and *RET51-mCherry* constructs and helpful discussion.

## Supplementary Data

**Supplementary Movie 1: Time lapse image of Ret51-Jip3 co-transport.** xMerged time lapse of Ret51-EGFP (green channel) and Jip3-mCherry (red channel) axonal transport acquired at 3 frames/sec. Retrograde transport of Ret51+ particles is marked by yellow arrowhead, anterograde transport of Ret51+ particle is marked by white arrowhead, and retrograde co-transport of Jip3+ Ret51+ particles is marked by magenta arrowhead. Scale bar = 5µm.

**Supplementary Figure 1:**
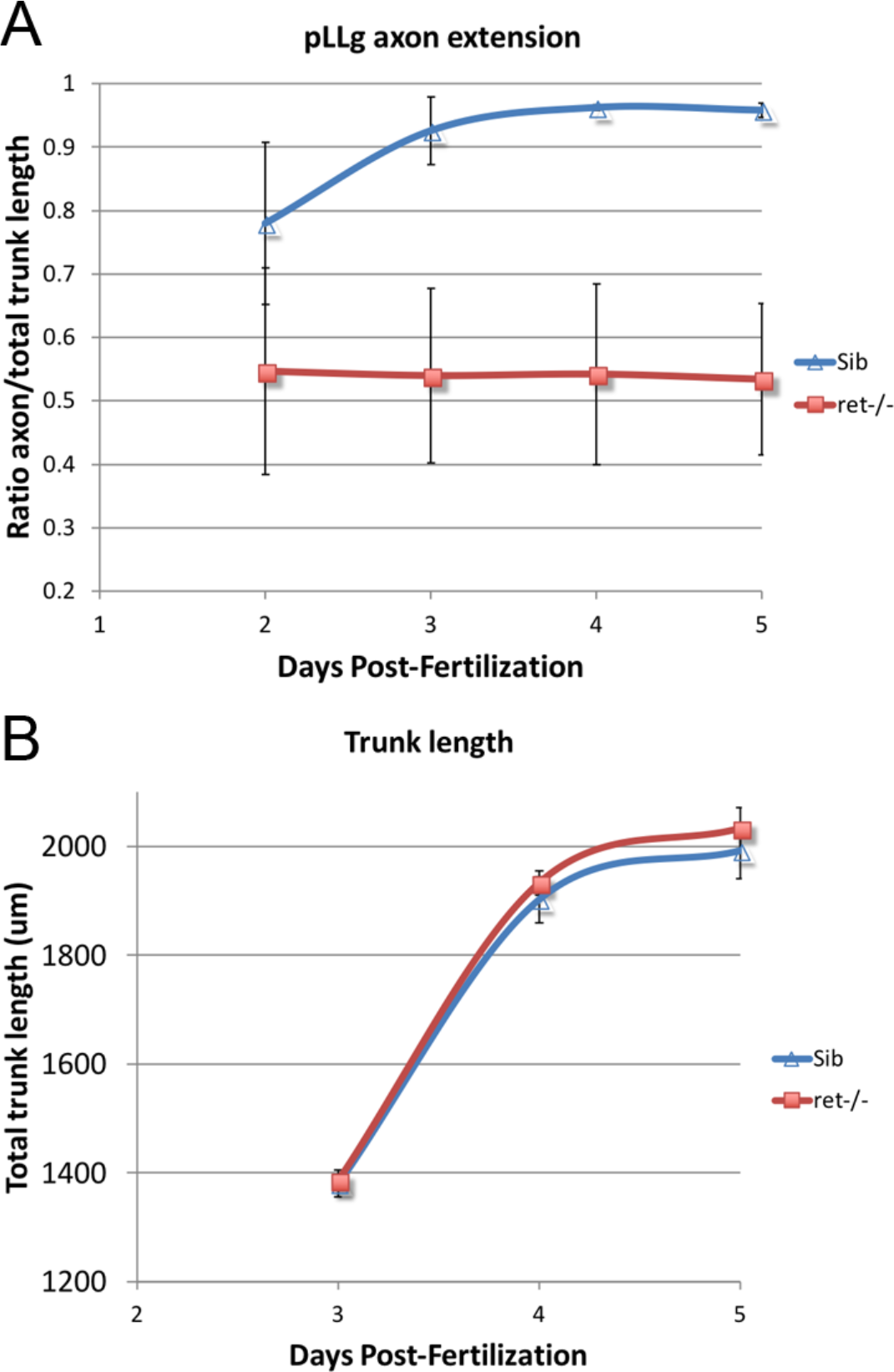
*ret^hu2846^* mutants do to display axon retraction or degeneration defects. (A) Ratio of length of extended pLLg axon in *neurod:EGFP* transgenic embryos over full trunk length from 2-5 dpf. *ret^hu2846^* mutants (n=6) display obvious axon truncation by 2 dpf compared to wild-type siblings (Sib, n=8), but mutant axons do not retract or degenerate over time (not significant by repeated measures ANOVA with post-hoc Bonferroni test for pairwise comparison; F(3,2)=0.043, p=0.985). (B) Measure of total trunk length sibling and *ret^hu2846^* mutant embryos from 3-5 dpf. No difference was found, indicating changing axon/trunk length ratio values are not altered by gross trunk morphology changes.

**Supplementary Figure 2:**
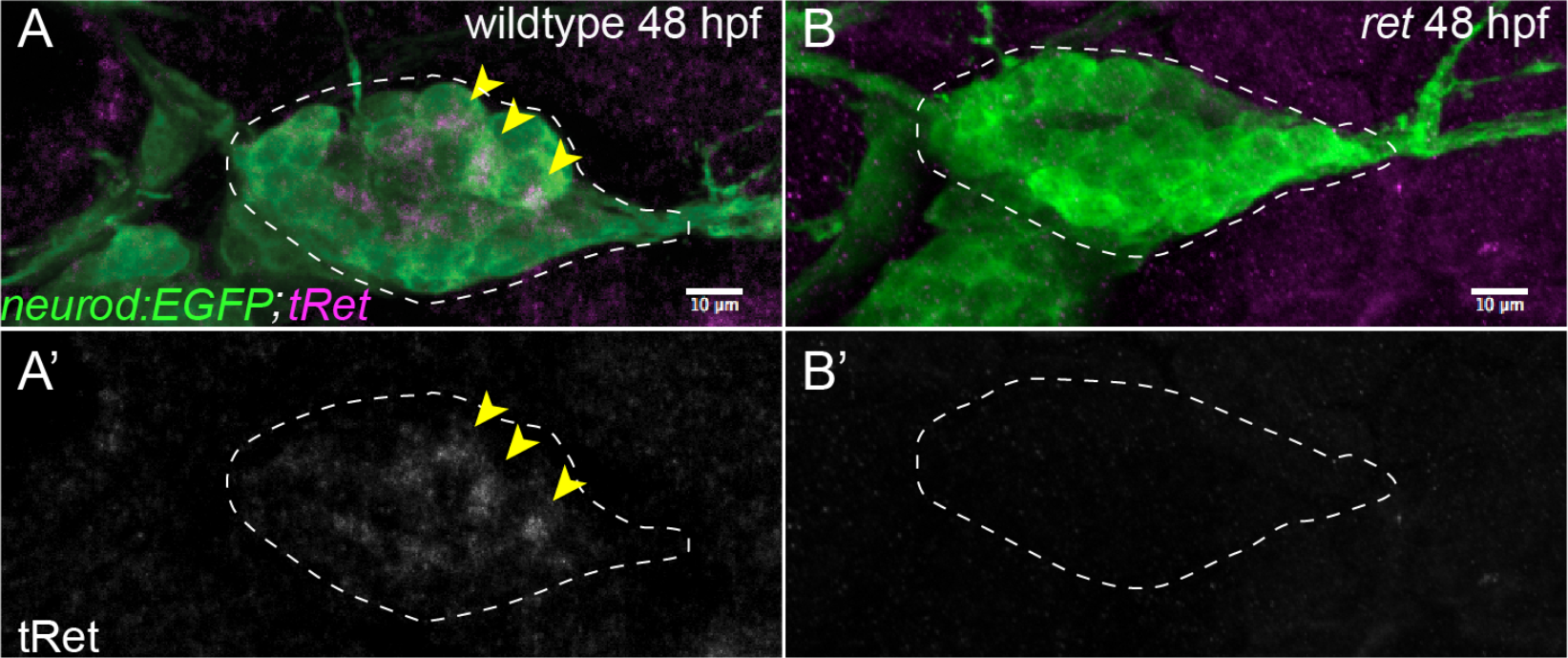
Anti-tRet antibody does not produce a detectable signal in *ret* mutants. Whole-mount immunolabeling of wild-type sibling (A) and *ret^hu2846^* mutant (B) with tRet antibody at 48 hpf. *neurod:EGFP* transgene marks all neurons in the pLL ganglion (outlined). tRet immunoreactivity (arrowheads) is present a subset of wild-type neurons but is completely absent from the *ret^hu2846^* mutant. N=8 wild-type sibs and N=6 mutants from two independent experiments.

**Supplementary Figure 3:**
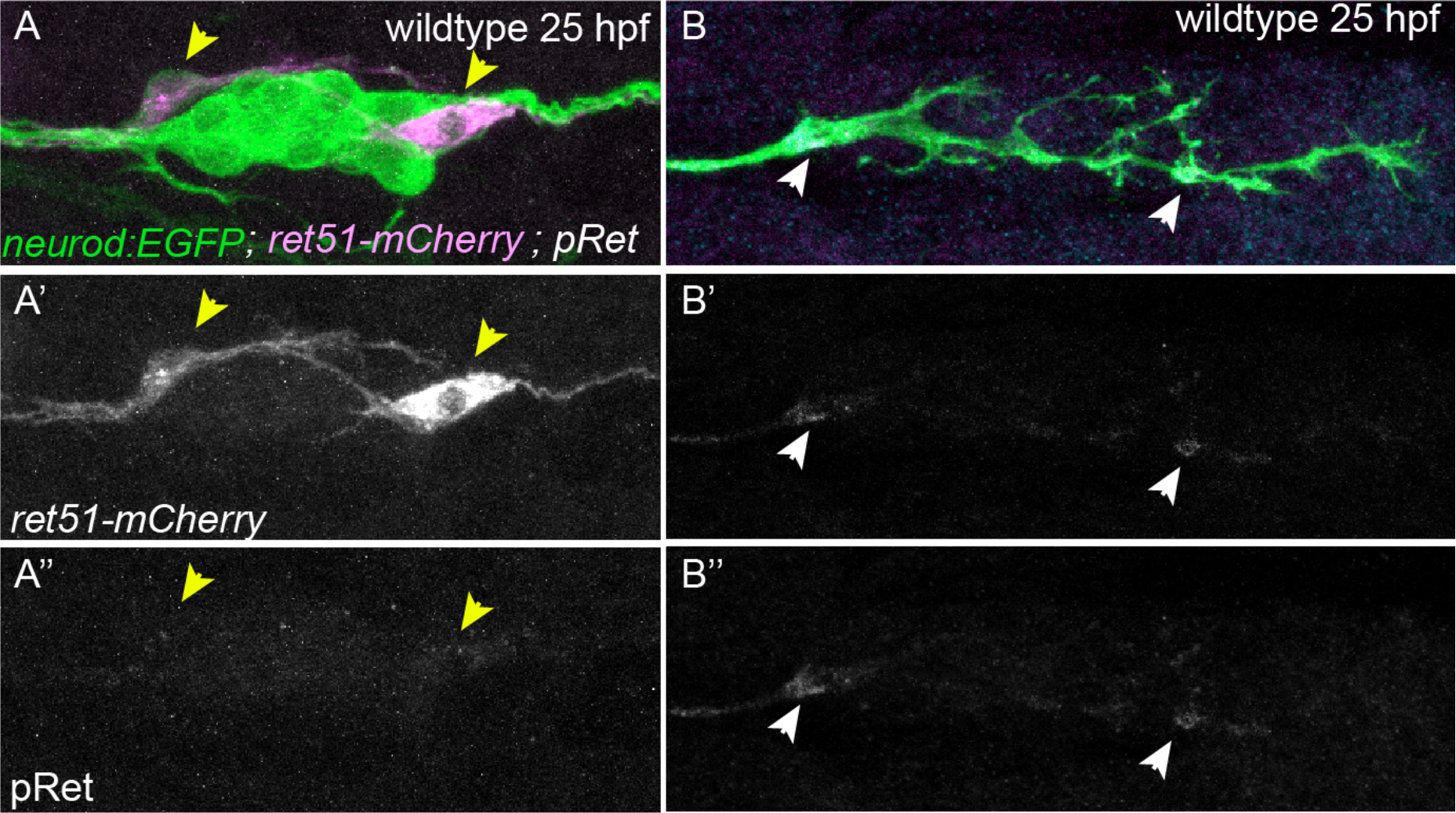
Anti-pRet antibody recognizes overexpressed RET51-mCherry fusion protein. TgBAC(*neurod:egfp*)-positive embryos were injected with 5 pg of *neurod5kb:RET51-mCherry* fusion plasmid and fixed at 25 hpf. Panels show immunolabeling of pLL ganglion (A) and pioneer growth cones (B) in wild-type embryo with pRet antibody. Note absence of the signal in the ganglion or cell bodies of neurons that express Ret51-mCherry (yellow arrowheads). However, pRet signal was reliably detected in growth cones (white arrowheads). N=12 cells from two independent experiments.

**Supplementary Figure 4:**
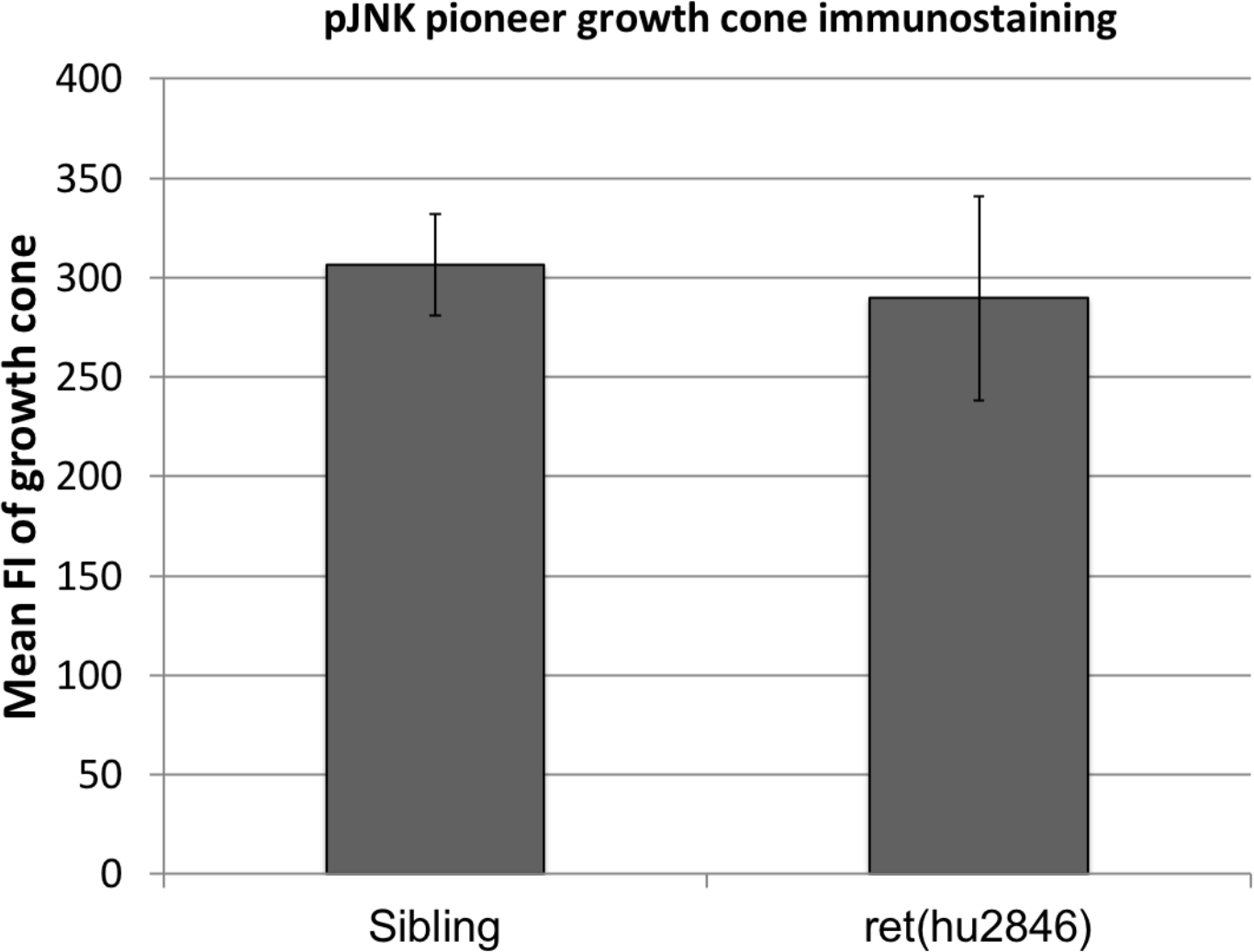
Anti-pJNK antibody immunostaining of pioneer growth cones. Mean fluorescence intensity measured for the most distal pLLg growth cones (visualized by *neurod:EGFP* transgene) was compared between sibling (306±26, n=14) and *ret^hu2846^* mutants (290±51, n=6) at 30 hpf, as not significantly different (*p*=0.78 by Mann-Whitney U test). Error bars represent S.E.M.

